# GraphBio: a shiny web app to easily perform popular visualization analysis for omics data

**DOI:** 10.1101/2022.02.28.482106

**Authors:** Tian-Xin Zhao, Ze-Lin Wang

**Affiliations:** Department of Pediatric Urology, Guangzhou Women and Children’s Medical Center, National Children’s Medical Center for South Central Region, Guangzhou Medical University, Guangzhou, Guangdong, China; Department of Pediatric Surgery, Guangzhou Institute of Pediatrics, Guangzhou Women and Children’s Medical Center, National Children’s Medical Center for South Central Region, Guangzhou Medical University, Guangzhou, Guangdong, China; Department of Bioinformatics, Shuzhi Biotech, LLC, Guangzhou, Guangdong, China

## Abstract

**Summary:** Here, we present GraphBio, a shiny web app to easily perform visualization analysis for omics data. GraphBio provides 15 popular visualization analysis methods, including heatmap, volcano plots, MA plots, network plots, dot plots, chord plots, pie plots, four quadrant diagrams, venn diagrams, cumulative distribution curves, PCA, survival analysis, ROC analysis, correlation analysis and text cluster analysis. It enables experimental biologists without programming skills to easily perform popular visualization analysis and get publication-ready figures.

**Availability and implementation:** GraphBio, as an online web application, is freely available at http://www.graphbio1.com/en/ (English version) and http://www.graphbio1.com/ (Chinese version).

## 1 Introduction

With the advance of high-throughput techniques (Goodwin, et al., 2016), more and more researchers start to depict molecular profiling in a systematic manner (Cancer Genome Atlas, 2012; Cancer Genome Atlas Research Network. Electronic address and Cancer Genome Atlas Research, 2017; Wang, et al., 2018; Yan, et al., 2015; Zhao, et al., 2022). Massive amount of omics data is produced and usually requires sophisticated visualization analysis. For example, gene expression studies frequently use heatmap, volcano plots and MA plots to characterize expression changes from thousands of genes (Conesa, et al., 2016). Moreover, PCA and correlation analysis are widely used to estimate similarity or dissimilarity between samples or groups. Despite these methods are popular in omics research, while they were usually published as R packages, such as ggplot2 (Wickham, 2016), pheatmap for drawing heatmap, GOplot for drawing chord plot of GO analysis results (Walter, et al., 2015), FactoMineR for performing PCA analysis (Lê, et al., 2008), pROC for drawing ROC curves (Robin, et al., 2011) et al, which require experimental biologists to have good programming skills. This makes a barrier for many experimental biologists without programming skills to use these methods.

To address this problem, we here developed an online web application called GraphBio using shiny framework in R software. GraphBio provides 15 popular visualization analysis methods, including heatmap, volcano plots, MA plots, network plots, dot plots, chord plots, pie plots, four quadrant diagrams, venn diagrams, cumulative distribution curves, PCA, survival analysis, ROC analysis, correlation analysis and text cluster analysis. It allows experimental biologists without programming skills to easily perform visualization analysis and get publication-ready plots in a short time.

## 2 Implementation

GraphBio was built using R package bs4Dash (Granjon, 2022), a Bootstrap 4 style-based Shiny dashboard web application framework, in RStudio and R software (version 4.0.3). All plots generated by GraphBio are entirely based on popular R packages, including ggplot2 (Wickham, 2016), pheatmap, survminer, pROC (Robin, et al., 2011), FactoMineR (Lê, et al., 2008), factoextra, GOplot (Walter, et al., 2015) and others. As an online web application, it is freely available at http://www.graphbio1.com/en/ (English version) and http://www.graphbio1.com/ (Chinese version) without login requirement. It can be accessed through any web browsers, such as Google Chrome, Microsoft Edge et al.

## 3 Results

GraphBio provides 15 visualization analysis modules, including heatmap, volcano plots, MA plots, network plots, dot plots, chord plots, pie plots, four quadrant diagrams, venn diagrams, cumulative distribution curves, PCA, survival analysis, ROC analysis, correlation analysis and text cluster analysis. Some representative visualization results are shown in Fig. 1. For each module, we provided “run example” button located at the sidebar panel to help users rapidly view the output style of GraphBio. We also provide “view example file” button to help users rapidly know the format of input files. Users can create a csv or comma-separated text file similar to the given example file and upload it to GraphBio by clicking “Browse” button in the sidebar panel. Once the file has been uploaded, the plot is automatically created with default parameters. Users can change default configurations through parameter settings in the sidebar panel. To avoid users being faced with too many parameters to know where to start, we only provide several necessary and easy-to-understand parameter choices, such as colors, font size et al. After the figure is completed, users can specify figure size and download it as a PDF file by clicking the “Download PDF Figure” button at the bottom of sidebar panel. The figures are saved in a vector format, thus ensuring the high quality of them. In addition, the figures can subsequently be easily edited using Adobe Illustrator software.

**Fig. 1.**
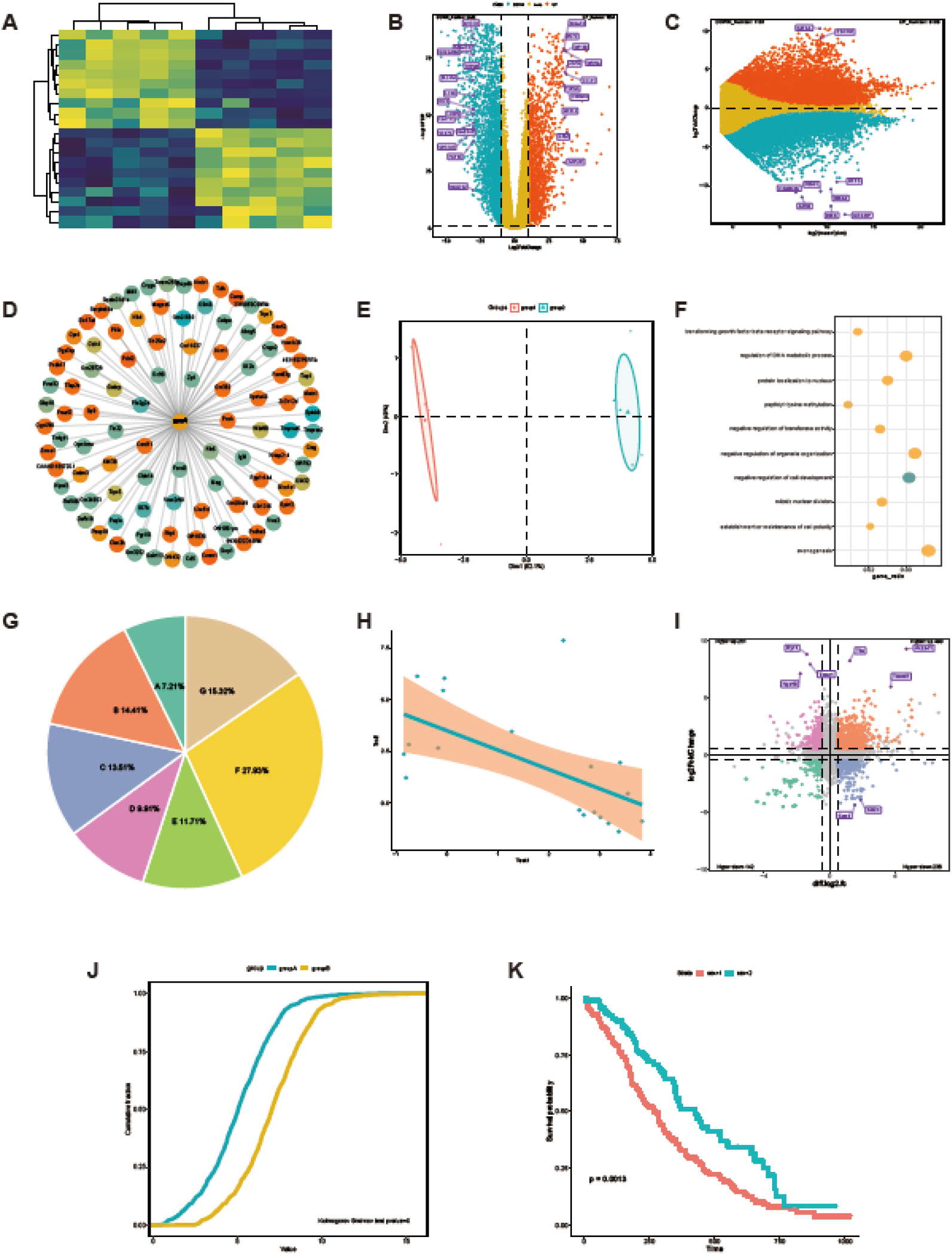
Some representative visualization results from GraphBio. (**A**) Gene expression heatmap. Rows represent genes and columns represent samples. Yellow represents up-regulation, while blue represents down-regulation. (**B**) Volcano plot shows significantly differentially expressed genes. X-axis represents log2-transformed fold changes, y-axis represents −log10(FDR). Red represents up-regulation, blue represents down-regulation, yellow represents genes which are not statistically significant. Some genes of interest are marked in purple. (**C**) MA plot shows significantly differentially expressed genes. X-axis represents mean expression values of genes, y-axis represents log2-transformed fold changes. Red represents up-regulation, blue represents down-regulation, yellow represents genes which are not statistically significant. Some genes of interest are marked in purple. (**D**) Network plot shows a group of expression-related genes for a target gene. The correlation values are calculated using Pearson correlation analysis. Red represents positive correlation, while blue represents negative correlation. (**E**) PCA analysis. (**F**) Dot plot shows some biological processes of interest. X-axis represents gene ratio. Point size represents gene numbers. Color represents significance. (**G**) Pie plot. (**H**) Pearson correlation analysis between two variables. (**I**) Four quadrant diagram shows the overlapped genes between RNA-seq and m6A-seq data. X-axis represents log2-transformed fold changes of m6A-seq data, y-axis represents log2-transformed fold changes of RNA-seq data. Significant genes are marked in four different colors. (**J**) Cumulative distribution curves. Kolmogorov-Smirnov test is used for comparing two samples. (**K**) Survival curves. Log-rank test is used for comparing two samples.

## 4 Conclusion

In this article, we introduce GraphBio, an online web application for omics data visualization. It includes 15 popular visualization analysis methods, such as heatmap, volcano plots, MA plots et al. Experimental biologists can easily perform online analysis and get publication-ready plots via access the website http://www.graphbio1.com/en/ (English version) or http://www.graphbio1.com/ (Chinese version) using any web browsers like Google Chrome, Microsoft Edge et al. In the future, we will continue to integrate more popular visualization analysis methods into GraphBio and provide more support to research community.

## Funding

This work was supported by the Guangdong Basic and Applied Basic Research Foundation (No. 2020A1515110796), the Research Fund for Clinical Doctor of Guangzhou Women and Children’s Medical Center, and the Research Fund from Shuzhi Biotech, LLC, Guangzhou.

## Conflict of Interest

none declared.

## Notes

### Competing Interest Statement

The authors have declared no competing interest.

